# RNA extension drives a stepwise displacement of an initiation-factor structural module in initial transcription

**DOI:** 10.1101/855544

**Authors:** Lingting Li, Vadim Molodtsov, Wei Lin, Richard H. Ebright, Yu Zhang

## Abstract

All organisms--bacteria, archaea, and eukaryotes--have a transcription initiation factor that contains a structural module that binds within the RNA polymerase (RNAP) active-center cleft and interacts with template-strand single-stranded DNA (ssDNA) in the immediate vicinity of the RNAP active center. This transcription-initiation-factor structural module pre-organizes template-strand ssDNA to engage the RNAP active center, thereby facilitating binding of initiating nucleotides and enabling transcription initiation from initiating mononucleotides. However, this transcription-initiation-factor structural module occupies the path of nascent RNA and thus presumably must be displaced before or during initial transcription. Here, we report four sets of crystal structures of bacterial initially transcribing complexes that demonstrate, and define details of, stepwise, RNA-extension-driven displacement of the “σ finger” of the bacterial transcription initiation factor σ. The structures reveal that--for both the primary σ factor and extracytoplasmic (ECF) σ factors, and for both 5’-triphosphate RNA and 5’-hydroxy RNA--the “σ finger” is displaced in stepwise fashion, progressively folding back upon itself, driven by collision with the RNA 5’-end, upon extension of nascent RNA from ∼5 nt to ∼10 nt.

**SIGNIFICANCE STATEMENT:** The “σ finger” of the bacterial initiation factor σ binds within the RNA polymerase active-center cleft and blocks the path of nascent RNA. It has been hypothesized that the σ finger must be displaced during initial transcription. By determining crystal structures defining successive steps in initial transcription, we demonstrate that the σ finger is displaced in stepwise fashion, driven by collision with the RNA 5’-end, as nascent RNA is extended from ∼5 nt to ∼10 nt during initial transcription, and we show that this is true for both the primary σ factor and alternate σ factors. Stepwise displacement of the σ finger can be conceptualized as stepwise compression of a “protein spring” that stores energy for subsequent breakage of protein-DNA and protein-protein interactions in promoter escape.

## INTRODUCTION

All cellular RNA polymerases (RNAPs)--bacterial, archaeal, and eukaryotic--require a transcription initiation factor, or a set of transcription initiation factors, to perform promoter-specific transcription initiation (refs. 1-4). In all promoter-specific transcription initiation complexes--bacterial, archaeal, and eukaryotic--a structural module of a transcription initiation factor enters the RNAP active-center cleft and interacts with template-strand single-stranded DNA (ssDNA) close to the RNAP active center (refs. 2, 5-27). This transcription-initiation-factor structural module pre-organizes template-strand ssDNA to adopt a helical conformation and to engage the RNAP active center, thereby facilitating binding of initiating nucleotides and enabling primer-independent transcription initiation from initiating mononucleotides (refs. 1-9, 11, 12, 16-19, 24, 27-29). However, this transcription-initiation-factor structural module occupies the path that will be occupied by nascent RNA during initial transcription and thus presumably must be displaced before or during initial transcription (refs. 1-3, 5-12, 17-18, 28, 30-33).

In bacterial transcription initiation complexes containing the primary σ factor and most alternative σ factors, the relevant structural module is the σ-factor “σ finger” (also referred to as σR3.2, σR3-σR4 linker, or the σR2-σR4 linker) (refs. 1, 2, 5-10, 14, 34). In bacterial transcription initiation complexes containing the alternative σ factor, σ^54^, the relevant structural module is the σ^54^ RII.3 region, which is structurally unrelated to the σ finger (refs. 13, 15). In archaeal RNAP-, eukaryotic RNAP-I-, eukaryotic RNAPII-, and eukaryotic RNAP-III-dependent transcription initiation complexes, the relevant structural modules are the TFB zinc ribbon and CSB, the Rrn7 zinc ribbon and B-reader, the TFIIB zinc ribbon and B-reader, and the Brf1 zinc ribbon, respectively, all of which are structurally related to each other but are structurally unrelated to the bacterial σ finger and σ^54^ RII.3 (refs. 16-18, 21-27).

It has been apparent for nearly two decades that the relevant structural module must be displaced before or during initial transcription (refs. 1-3, 5-12, 17-18, 28-33). Changes in protein-DNA photo-crosslinking suggestive of displacement have been reported (ref. 1), and changes in profiles of abortive initiation and initial-transcription pausing upon mutation of the relevant structural module, have been reported (refs. 1, 9, 24, 28-29, 31-33, 35-36). However, the displacement has not been observed directly. As a result, it has remained unclear whether the displacement occurs in a single step or in multiple steps, it has remained unclear where the displaced residues move, and it has remained unclear what drives the displacement.

Here, we present four sets of crystal structures of bacterial initially transcribing complexes that demonstrate--and define structural and mechanistic details of--displacement of the σ finger during initial transcription. The first and second sets are structures of initial transcribing complexes comprising *Thermus thermophilus* (*Tth*) RNAP, the *Tth* primary σ factor, σ^A^, promoter DNA, and either 5’-hydroxyl-containing RNAs 2, 3, 4, 5, 6, and 7 nt in length (σ^A^-RPitc2, σ^A^-RPitc3, σ^A^-RPitc4, σ^A^-RPitc5, σ^A^-RPitc6, and σ^A^-RPitc7; Fig. 1) or comprising *Tth* RNAP, *Tth σ*^A^, promoter DNA, and 5’-triphosphate-containing RNAs, 2, 4, 5, and 6 nt in length (σ^A^-RPitc2-PPP, σ^A^-RPitc4-PPP, σ^A^-RPitc5-PPP, and σ^A^-RPitc6-PPP; Fig. 2). These sets correspond to, respectively, six successive intermediates in primary-σ-factor-dependent, primer-dependent transcription initiation (5’-hydroxyl-containing RNA; ref. 2), and four successive intermediates in primary-σ-factor-dependent, primer-independent transcription initiation (5-triphosphate-containing RNA; ref. 2). The third and fourth sets are structures of initial transcribing complexes comprising *Mycobacterium tuberculosis* (*Mtb*) RNAP, *Mtb* extracytoplasmic-function (ECF) σ factor σ^H^, promoter DNA, and 5’-hydroxyl-containing RNA oligomers 5, 6, 7, 9, and 10 nt in length (σ^H^-RPitc5, σ^H^-RPitc6, σ^H^-RPitc7, σ^H^-RPitc9, and σ^H^-RPitc10; Fig. 3) and comprising *Mtb* RNAP, *Mtb* ECF σ factor σ^L^, promoter DNA, and 5’-hydroxyl-containing RNA oligomers 5, 6, and 9 nt in length (σ^L^-RPitc5, σ^L^-RPitc6, and σ^L^-RPitc9; Fig. 4). These sets correspond to, respectively, five intermediates in alternative-σ-factor-dependent transcription initiation with ECF σ factor σ^H^, and three intermediates in alternative-σ-factor σ^L^-factor-dependent transcription initiation with ECF σ factor σ^L^. Taken together, the structures establish that--for both the primary σ factor and ECF σ factors, and for both 5’-hydroxyl RNA and 5’-triphosphate RNA--the “σ finger” is displaced in stepwise fashion, progressively folding back upon itself, driven by collision with the RNA 5’-end, upon extension of nascent RNA from 5 nt to ∼10 nt.

**Figure 1.**
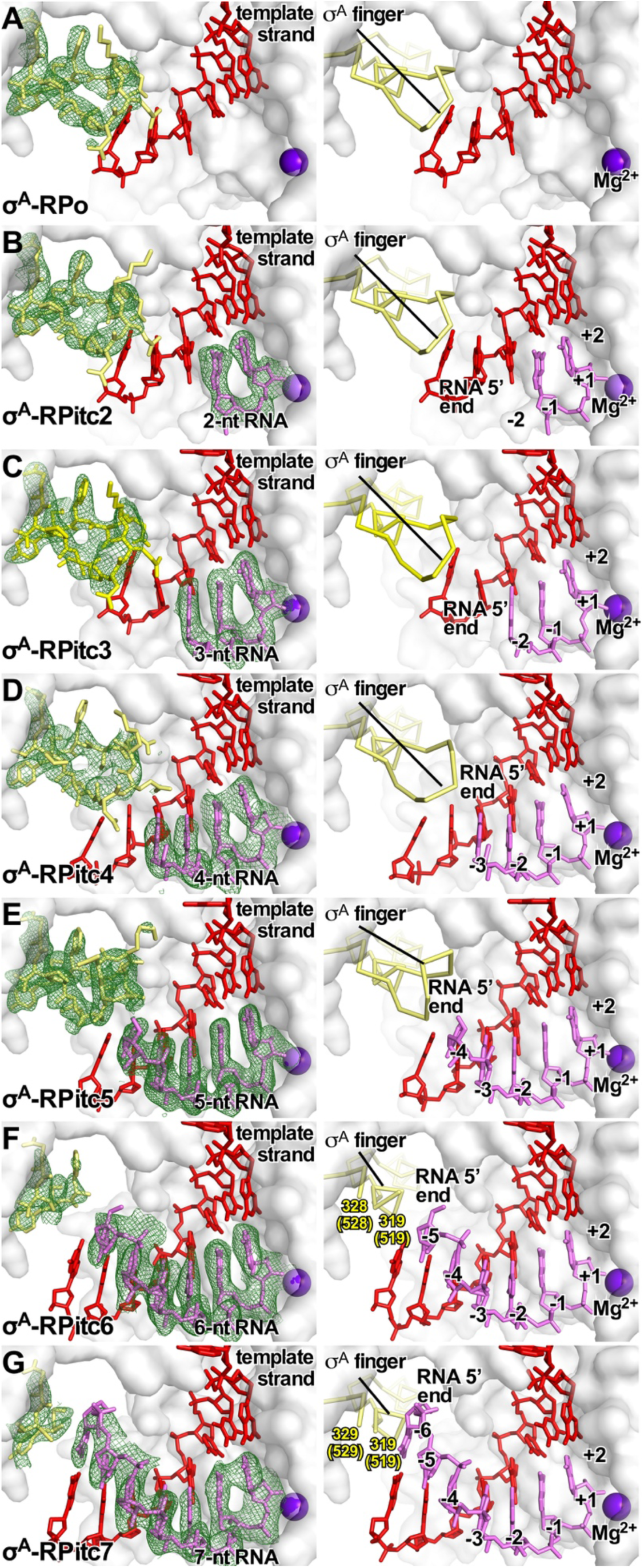
Stepwise, RNA-driven displacement of the σ finger: primary-σ factor, 5’-hydroxyl RNA. Crystal structures of **(A)** σ^A^-RPo (PDB 4G7H; ref. *5*), **(B)** σ^A^-RPitc2 (PDB 4G7O; ref. *5*), **(C)** σ^A^-RPitc3 (this work; Table S1), **(D)** σ^A^-RPitc4 (this work; Table S1), **(E)** σ^A^-RPitc5 (this work; Table S1), **(E)** σ^A^-RPitc6 (this work; Table S1), and **(E)** σ^A^-RPitc7 (this work; Table S1). Left subpanels, electron-density map and atomic model. Right subpanels, atomic model. Green mesh, simulated-annealing Fo-Fc electron-density map contoured at 2.5 σ; gray surface, RNAP β’ subunit; yellow sticks, σ finger; red sticks, template-strand DNA; pink sticks, RNA; purple sphere, catalytic Mg^2+^ ion.

**Figure 2.**
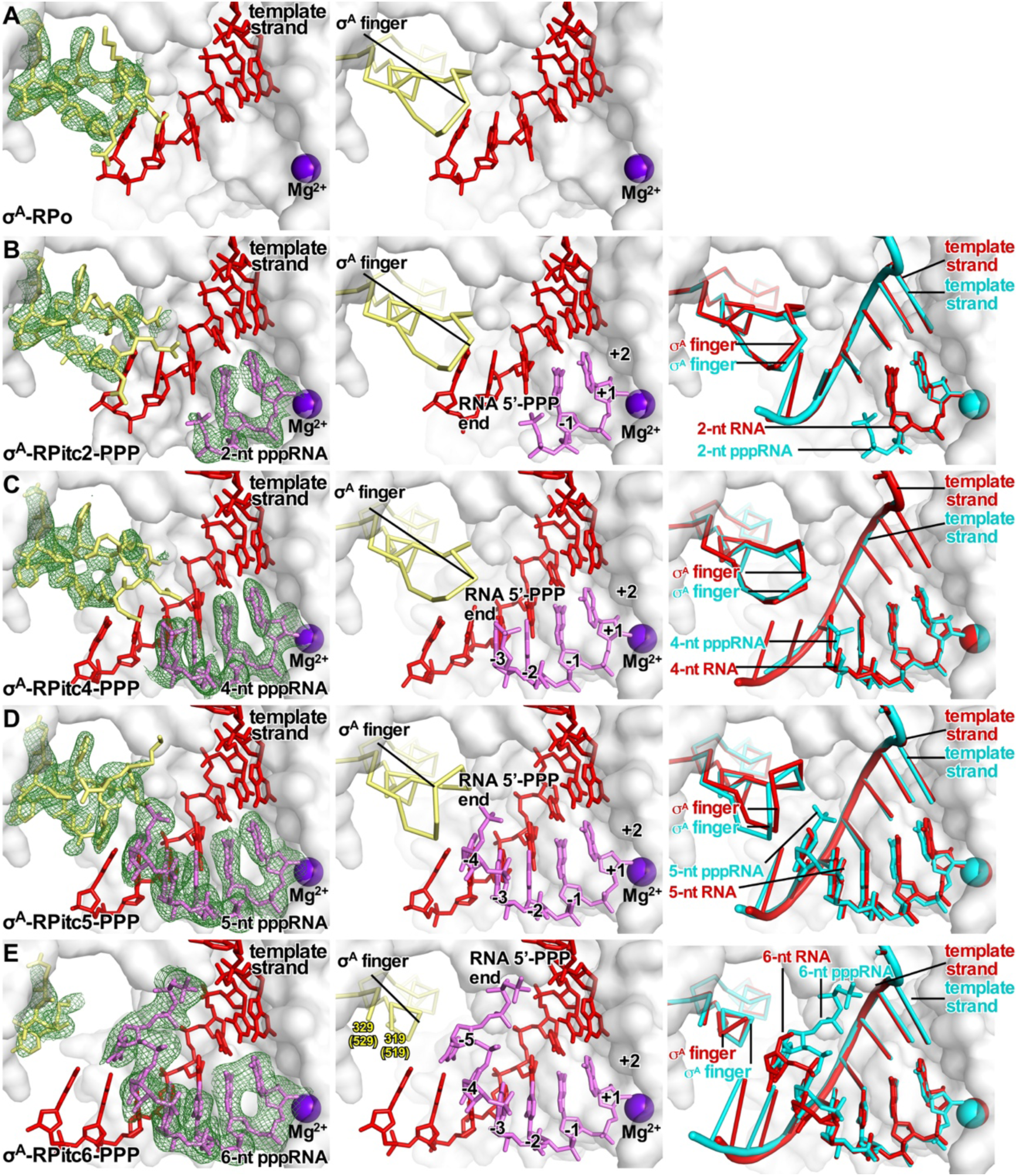
Stepwise, RNA-driven displacement of the σ finger: primary σ factor, 5’-triphosphate RNA. Crystal structures of **(A)** σ^A^-RPo (PDB 4G7H; ref. *5*), **(B)** σ^A^-RPitc2-PPP (this work; Table S2), **(C)** σ^A^-RPitc4-PPP (this work; Table S2), **(D)** σ^A^-RPitc5-PPP (this work; Table S2), **(E)** σ^A^-RPitc6-PPP (this work; Table S2). Left subpanels, electron-density map and atomic model (colors as in Fig 1). Center subpanels, atomic model (colors as in Fig. 1). Right subpanel, Superimposition of structures of corresponding complexes with 5’-hydroxyl RNA (cyan and gray) and 5’-triphosphate RNA (red and gray).

**Figure 3.**
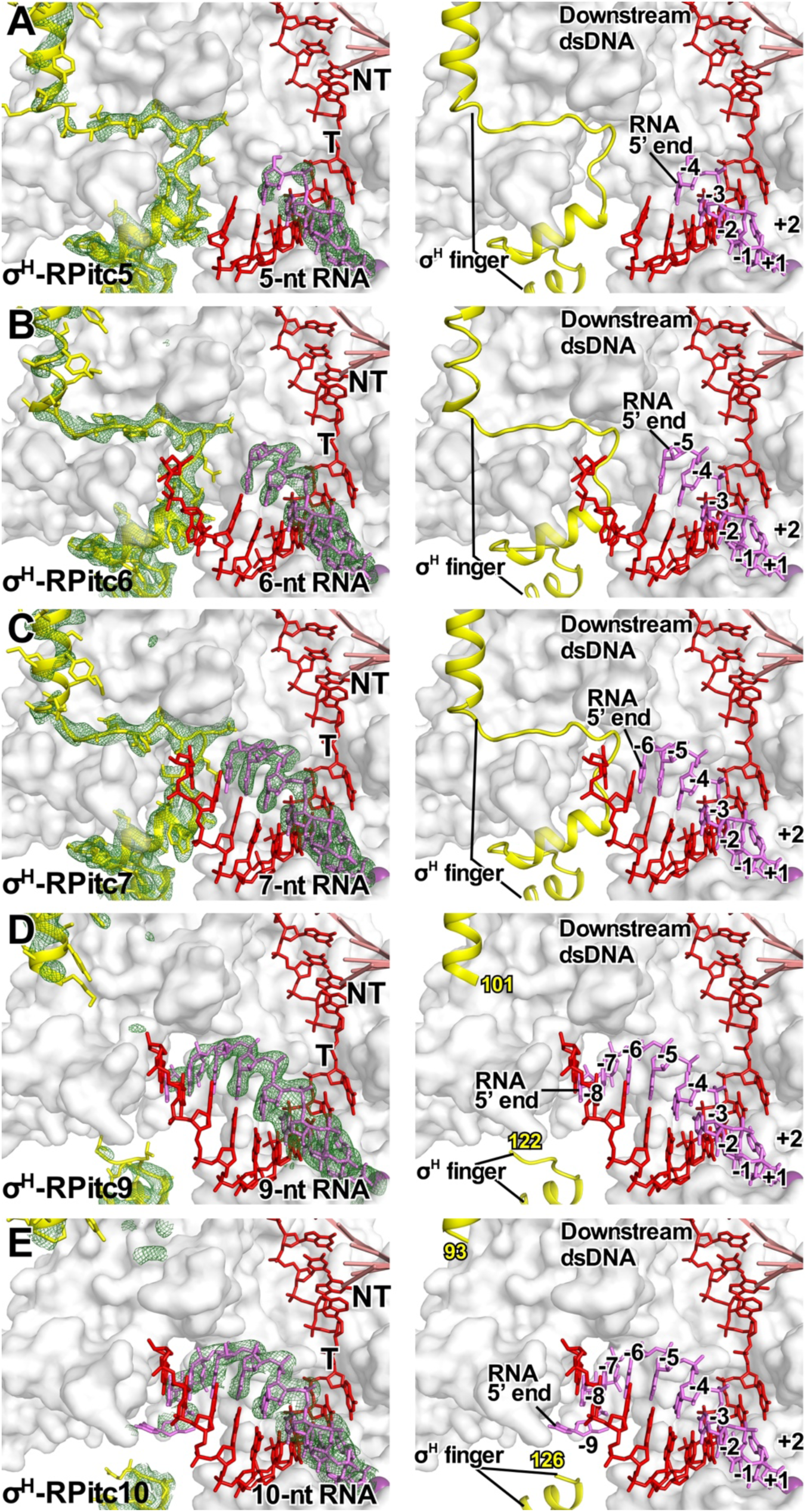
Stepwise, RNA-driven displacement of the σ finger: ECF σ factor σ^H^. Crystal structures of **(A)** σ^H^-RPitc5 (this work; Table S3), **(B)** σ^H^-RPitc6 (PDB 6JCX; ref. 6), **(C)** σ^H^-RPitc7 (this work; Table S3), **(D)** σ^H^-RPitc9 (this work; Table S3), and **(E)** σ^H^-RPitc10 (this work; Table S3). Colors as in Fig. 1.

**Figure 4.**
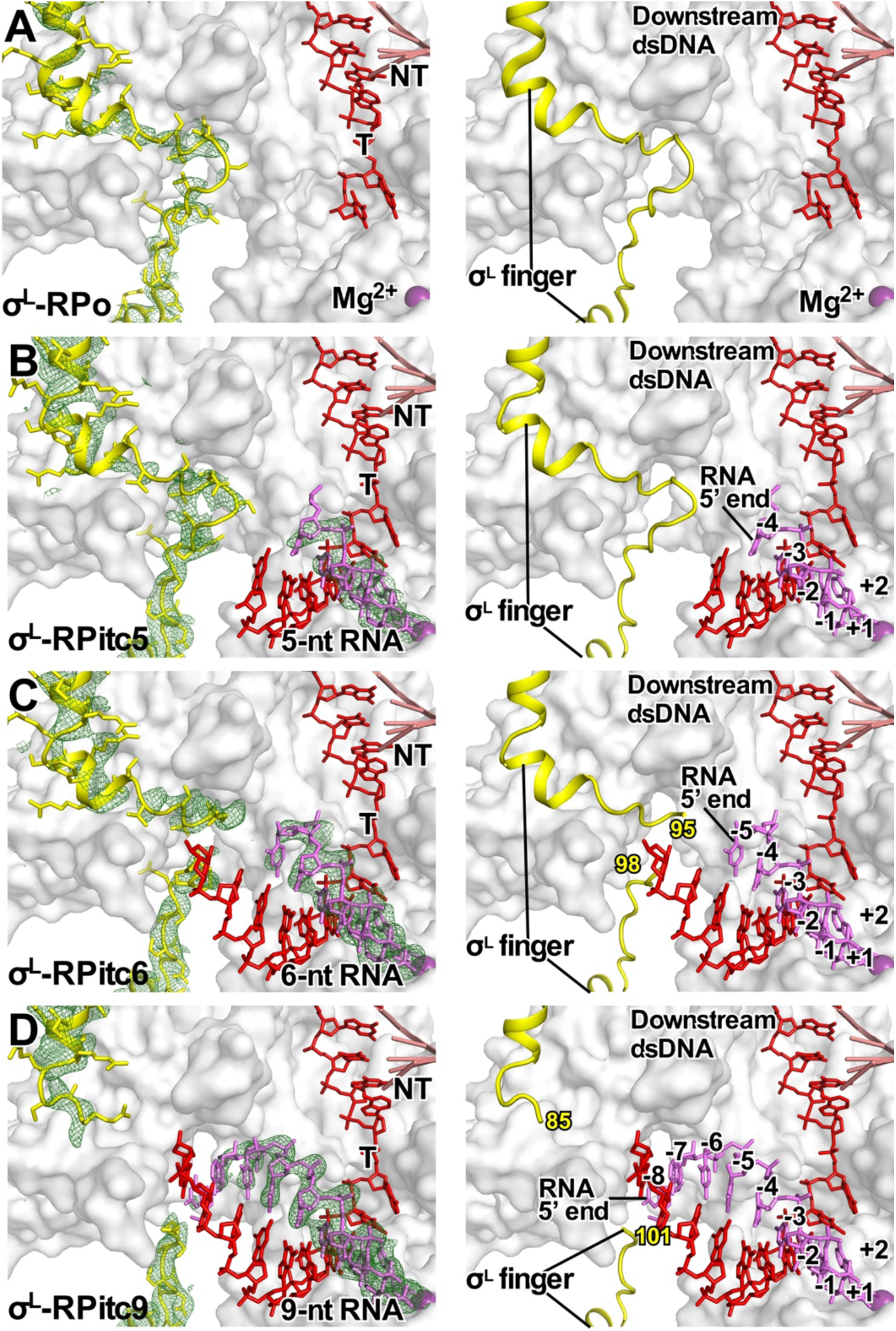
Stepwise, RNA-driven displacement of the σ finger: ECF σ factor σ^L^. Crystal structures of **(A)** σ^L^-RPo (PDB 6DVE; ref. 7), **(B)** σ^L^-RPitc5, (this work; Table S4; 7), **(C)** σ^L^-RPitc6 (this work, Table S4), and **(D)** σ^L^-RPitc9 (this work, Table S4). Colors as in Fig. 1.

## RESULTS

### Stepwise, RNA-driven displacement of the σ finger: primary σ, 5’-hydroxyl RNA

The primary σ factor, responsible for transcription initiation at most promoters under most conditions, is σ^A^ (σ^70^ in *E. coli*; ref. 1).

Transcription initiation can proceed through either a primer-dependent pathway or a primer-independent pathway (ref. 2). In primer-dependent transcription initiation, the initiating entity is a short RNA oligomer, typically a 5’-hydroxyl-containing short RNA oligomer, which yields a nascent RNA product containing a 5’-hydroxyl end.

To define the structural and mechanistic details of displacement of the σ finger during primary-σ-factor-dependent, primer-dependent transcription initiation, we determined crystal structures of initial transcribing complexes comprising *Tth* RNAP, the *Tth* primary σ factor, σ^A^, promoter DNA, and 5’-hydroxyl-containing RNAs 3, 4, 5, 6, and 7 nt in length (*Tth σ*^A^-RPitc3, σ^A^-RPitc4, σ^A^-RPitc5, σ^A^-RPitc6, and σ^A^-RPitc7; Figs. 1C-G and S1A; Table S1). In each case, the crystal structure shows a post-translocated transcription initiation complex (ref. 2), in which the RNA oligomer forms a base-paired RNA-DNA hybrid with template-strand DNA and is positioned such that the RNA 3’ nucleotide is located in the RNAP active-center product site (P site), in an orientation that would allow binding of, and base stacking by, an NTP substrate in the RNAP active-center addition site (A site) (Figs. 1 and S1A). Thus, in *Tth σ*^A^-RPitc3, σ^A^-RPitc4, σ^A^-RP itc5, σ^A^-RPitc6, and σ^A^-RPitc7, the RNA oligomers are base paired with template-strand DNA positions −2 to +1, −3 to +1, −4 to +1, −5 to +1, and −6 to +1, respectively (Fig. 1C-G). Together with the previously reported crystal structures of *Tth σ*^A^-RPo and σ^A^-RPitc2 (PDB 4G7H and PDB 4G7O; ref. 5), the new crystal structures constitute a series of structures in which the position of the RNA 5’-end relative to RNAP varies in single-nucleotide increments, and in which the position of RNA 3’-end relative to RNAP remains constant, as in stepwise RNA extension in initial transcription (Fig. 1).

In the crystal structure of *Tth σ*^A^-RPitc3, the σ finger adopts the same conformation as in the previously reported structures of *Tth σ*^A^-RPo and σ^A^-RPitc2 (Figs. 1A-C; ref. 5). Residues D514, D516, S517, and F522 of the σ^A^ finger (σ^A^ residues numbered as in *E. coli σ*^70^ here and hereafter) make H-bonded and van der Waals interactions with the template-strand DNA nucleotides at positions −3, −4, and −5, pre-organizing the template strand to adopt a helical conformation and to engage the RNAP active center (Fig. S2C).

In the crystal structure of *Tth σ*^A^-RPitc4, the σ finger adopts the same conformation as in the structures of *Tth σ*^A^-RPo, σ^A^-RPitc2, and σ^A^-RPitc3 (Figs. 1A-D), and residues D514, S517, and F522 of the σ^A^ finger again make H-bonded and van der Waals interactions with the template-strand DNA nucleotides at positions −3, −4, and −5, pre-organizing the template strand to adopt a helical conformation and to engage the RNAP active center (Fig. S2D). However, in RPitc4, residues D513 and D514 of the σ^A^ finger now also make H-bonded and van der Waals interactions with RNA, contacting RNA nucleotides at positions −3 and −4 in a manner that likely stabilizes the RNA-DNA hybrid in σ^A^-RPitc4.

In the crystal structure of *Tth σ*^A^-RPitc5, the σ finger adopts a different conformation as compared to the structures of *Tth σ*^A^-RPo, σ^A^-RPitc2, σ^A^-RPitc3, and σ^A^-RPitc4 (Figs. 1A-E), and this difference in conformation coincides with, and likely is driven by, the movement of the 5’ end of RNA into the position previously occupied by the tip of the σ finger and resulting steric collision between the RNA and the σ finger (Figs. 1D-E). In σ^A^-RPitc5, as compared to in σ^A^-RPo and σ^A^-RPitc2 through σ^A^-RPitc4, the tip of the σ^A^ finger--σ residues 513, 514, and 515--folds back over the rest of the σ finger, resulting in the displacement, away from the RNAP active center, of the Cα atoms of residues 513, 514, and 515 by 3 Å, 6 Å, and 3 Å, respectively (Figs. 1D-E). The change in conformation can be best understood as the folding of the loop of the σ-finger β-hairpin back over the strands of the σ-finger β-hairpin (Figs. 1D-E). In this new conformation, the σ finger makes new interactions with the RNA-DNA hybrid, with σ residues I511 and F522 making van del Waals interactions with template-strand DNA at position −5, and with σ residues D513 and D516 making H-bonds with RNA at position −4.

In the crystal structures of *Tth σ*^A^-RPitc6 and σ^A^-RPitc7, the σ finger again adopts different conformations, and the differences in conformation again coincide with, and likely are driven by, movement of the 5’ end of RNA into the positions previously occupied by the tip of the σ finger and resulting steric collision between the RNA and the σ finger (Figs. 1F-G). In the structure of σ^A^-RPitc6, eight residues of the σ^A^ finger--σ residues 520-527--exhibit segmental disorder, indicating that this eight-residue segment is flexible and exists as an ensemble of multiple conformations (Fig. 1F). In the structure of σ^A^-RPitc7, nine residues of the σ^A^ finger--σ residues 520-528--exhibit segmental disorder, indicating that this nine-residue segment is flexible and exists as an ensemble of multiple conformations (Fig. 1G). We interpret the changes in conformation from σ^A^-RPitc5 to σ^A^-RPitc6 to σ^A^-RPitc7 as successive further displacements of the loop of the σ-finger β-hairpin--involving displacement of first an eight-residue segment and then a nine-residue segment--back over the strands of the σ-finger β-hairpin, accompanied by increased flexibility of the displaced segments (Figs. 1E-G).

Taken together, the crystal structures of *Tth σ*^A^-RPo, σ^A^-RPitc2, σ^A^-RPitc3, σ^A^-RPitc4, σ^A^-RPitc5, σ^A^-RPitc6, and σ^A^-RPitc7 demonstrate that nascent RNA with a 5’-hydroxyl end displaces the σ finger of the primary σ factor in a stepwise fashion, starting upon extension of nascent RNA to a length of 5 nt and continuing upon further extension through a length of at least 7 nt (Fig. 1).

### Stepwise, RNA-driven displacement of the σ finger: primary σ factor, 5’-triphosphate RNA

As noted above, transcription initiation can proceed through either a primer-dependent pathway or a primer-independent pathway (ref. 2). In primer-independent transcription initiation, the initiating entity is a mono-nucleotide, typically a nucleoside triphosphate (NTP), which yields a nascent RNA product containing a 5’-triphosphate end. As compared to the 5’-hydroxyl end typically generated in primer-dependent initiation, the 5’-triphosphate end typically generated in primer-independent initiation has greater charge (−4 charge vs. 0 charge) and greater steric bulk (∼160 Å^3^ solvent-accessible volume vs. ∼7 Å^3^ solvent accessible-volume). In an RNA-extension-driven displacement mechanism, the additional charge and steric bulk in principle could change the length-dependence or, even the pathway, of initiation-factor displacement. To assess whether the additional charge and steric bulk in fact change the length-dependence or pathway of initiation-factor displacement, we performed an analysis analogous to that in the preceding section, but assessing nascent RNA products containing 5’-triphosphate ends.

We determined crystal structures of initial transcribing complexes comprising *Tth* RNAP, the *Tth* primary σ factor, σ^A^, promoter DNA, and 5’-triphosphate-containing RNAs 2, 4, 5, and 6 nt in length (*Tth σ*^A^-RPitc2-PPP, σ^A^-RPitc4-PPP, σ^A^-RPitc5-PPP, and σ^A^-RPitc6-PPP; Figs. 2B-E and S1B; Table S2). In each case, the crystal structure shows a post-translocated transcription initiation complex, in which the RNA oligomer forms a base-paired RNA-DNA hybrid with template-strand DNA and is positioned such that the RNA 3’ nucleotide is located in the RNAP active-center P site, in an orientation that would allow binding of, and base stacking by, an NTP substrate in the RNAP active-center A site. Thus, in *Tth σ*^A^-RPitc2-PPP, σ^A^-RPitc4-PPP, σ^A^-RP itc5-PPP, and σ^A^-RPitc6-PPP, the RNA oligomers are base paired with template-strand DNA positions −1 to +1, −3 to +1, −4 to +1, and −5 to +1, respectively (Fig. 2B-E). Together with the previously reported crystal structure of *Tth σ*^A^-RPo (PDB 4G7H; refs. 5, 37), the new crystal structures constitute a nearly complete series of structures in which the position of the RNA 5’-end relative to RNAP varies in single-nucleotide increments, and in which the position of RNA 3’-end relative to RNAP remains constant, as in stepwise RNA extension in initial transcription (Fig. 2). Efforts were made to obtain corresponding crystal structures of complexes containing 5’-triphosphate-containing RNAs 3 nt and 7 nt in length, but these efforts were unsuccessful, instead yielding a structure of a fractionally translocated complex (refs. 38, 39) and a structure with a disordered RNA 5’-end, respectively.

In the crystal structure of *Tth σ*^A^-RPitc2-PPP, the σ finger adopts the same conformation as in the structures of *Tth σ*^A^-RPo (PDB 4G7H; Figs. 2A-B). As in RPo, residues D514, D516, S517, and F522 of the σ^A^ finger make H-bonded and van der Waals interactions with the template-strand DNA nucleotides at positions-3 and −4 in a manner that pre-organizes the template strand to adopt a helical conformation and to engage the RNAP active center (Figs. S3A-B). The RNAP β-subunit residue R687 (RNAP residues numbered as in *E. coli* RNAP here and hereafter) makes an H-bonded interaction with the α phosphate of the RNA 5’-triphosphate; and the template-strand DNA nucleotide at positions −2 makes an H-bonded interaction with the γ phosphate of the RNA 5’-triphosphate.

In the crystal structure of *Tth σ*^A^-RPitc4-PPP, the σ finger adopts the same conformation as in the structures of *Tth σ*^A^-RPo and σ^A^-RPitc2-PPP (Figs. 2A-C). Essentially as in σ^A^-RPo and σ^A^-RPitc2-PPP, residues D514, I511, and F522 of the σ^A^ finger make van der Waals interactions with the template-strand DNA nucleotides at positions −3, −4, and −5 in a manner that pre-organizes the template strand to adopt a helical conformation and to engage the RNAP active center (Figs. 2A-C and S3A-C); and, as in the structure of *Tth σ*^A^-RPitc4, residue D514 of the σ^A^ finger makes H-bonded and van der Waals interactions with RNA, contacting RNA nucleotides at positions −2 and −3 in a manner that likely stabilizes the RNA-DNA hybrid. RNAP β–subunit residue R540 makes a salt bridge with the α phosphate of the RNA 5’-triphosphate, and RNAP β-subunit residue N568 makes an H-bonded interaction with the α phosphate of the RNA 5’-triphosphate. The β and γ phosphates of the RNA 5’-triphosphate are disordered, suggesting that the β and γ phosphates do not adopt a defined conformation and do not make defined interactions (Fig. 2A). We point out that Steitz and co-workers have reported close proximity, and likely contact, between the σ finger and the RNA 5’-end in a low-resolution (∼5.5 Å resolution) crystal structure of an initial transcribing complex containing a 5’-triphosphate-containing 5 nt RNA product in a pre-translocated state (ref. 12). The position of the RNA 5’ end in our structure of σ^A^-RPitc4-PPP in a post-translocated-state corresponds to the position of the RNA 5’ end in their structure, and, in our structure, contacts between the σ finger and the RNA 5’ end are resolved and defined.

In the crystal structure of *Tth σ*^A^-RPitc5-PPP, the σ finger adopts a conformation that is different from the conformation in the structures of *Tth* RPo, *Tth* RPitc2-PPP, and RPitc4-PPP (Figs. 2A-D) and that is similar to the conformation in the structure of *Tth σ*^A^-RPitc5 (Figs. 1E and 2D). As described above for the difference in conformation between σ^A^-RPitc4 and σ^A^-RPitc5, the difference in conformation between σ^A^-RPitc4-PPP and σ^A^-RPitc5-PPP coincides with, and likely is driven by, movement of the 5’ end of RNA into the position previously occupied by the tip of the σ finger and resulting steric collision between the RNA and the σ finger (Figs. 1D-E and 2C-D). The loop of the σ-finger β-hairpin folds back over the strands of the σ-finger β-hairpin (Figs. 2D), and, in this new conformation, the σ finger makes new interactions with the RNA-DNA hybrid, with σ residues I511 and F522 making van del Waals interactions with template-strand DNA at position −5, and with σ residues D516 making H-bonds with RNA at position −4 (Figs. 2D). RNAP β-subunit residue R540 makes a salt bridge with the α-phosphate of the RNA 5’-triphosphate. The β and γ phosphates of the RNA 5’-triphosphate are disordered, suggesting that the β and γ phosphates do not adopt a defined conformation and do not make defined interactions (Fig. 2C). We note that the negatively charged RNA 5’-triphosphate (−4 charge) is located close to--∼3 Å--the negatively charged tip of the σ^A^ finger (−3 charge), and that this proximity between negatively charged groups must result in electrostatic repulsion. This electrostatic repulsion does not manifest itself in as a difference in conformation between σ^A^-RPitc5-PPPand *Tth σ*^A^-RPitc5 (Figs. 1E and 2D), but could manifest itself as a difference in kinetics of RNA extension for 5’-triphosphate-containing nascent RNA products vs. 5’-hydoxyl-containing nascent RNA products, possibly accounting for previously observed differences in abortive-product distributions (ref. 2) and differences in susceptibilities to initial-transcription pausing (ref. 33) for 5’-triphosphate-containing nascent RNA products vs. 5’-hydoxyl-containing nascent RNA products. We point out that Murakami and co-workers have reported disorder of part of the σ finger in a crystal structure of an initial transcribing complex containing a 5’-triphosphate-containing 6 nt RNA product in a pre-translocated state (ref. 11). The position of the RNA 5’ end in our structure of σ^A^-RPitc5-PPP in a post-translocated-state corresponds to the position of the RNA 5’ end in their structure, but, in our structure, the conformation of the full σ finger is defined.

In the crystal structure of *Tth σ*^A^-RPitc6-PPP, the σ finger adopts a conformation that is even more different from the conformation in the structure of *Tth* RPitc4-PPP (Figs. 2C-E) and that is similar to the conformation in the structure of *Tth σ*^A^-RPitc6 (Figs. 1F and 2E). In the structure of σ^A^-RPitc6-PPP, nine residues of the σ^A^ finger--σ residues 520-528--exhibit segmental disorder, indicating that this nine-residue segment is flexible and exists as an ensemble of multiple conformations (Fig. 2E). We interpret the successive changes in conformation from σ^A^-RPitc4-PPP to σ^A^-RPitc5-PPP to σ^A^-RPitc6-PPP as successive displacements of the loop of the σ-finger β-hairpin back over the strands of the σ-finger β-hairpin, accompanied by increases in flexibility of the displaced segment (Fig. 2C-E). In the structure of σ^A^-RPitc6-PPP, RNAP β’-subunit residue K334 makes a salt bridge with the β phosphate of the RNA 5’-triphosphate. The α and γ phosphates of the RNA 5’-triphosphate are ordered but make no interactions with RNAP and σ.

Taken together, the crystal structures of *Tth σ*^A^-RPo, σ^A^-RPitc2-PPP, σ^A^-RPitc4-PPP, σ^A^-RPitc5-PPP, and σ^A^-RPitc6-PPP demonstrate that nascent RNA with a 5’-triphosphate end--like nascent RNA with a 5’-hydroxyl end--displaces the σ finger of the primary σ factor in a stepwise fashion, starting upon extension of nascent RNA to a length of 5 nt and continuing upon further extension through a length of at least 6 nt (Fig. 2).

### Stepwise, RNA-driven displacement of the σ finger: ECF σ factor σ^H^

The largest and most functionally diverse class of alternative σ factors is the extracytoplasmic function) ECF class (refs. 1, 40-43). ECF σ factors consist of a structural module (σ_2_) that recognizes a promoter −10 element, a structural module (σ_4_) that recognizes a promoter −35 element, and a σ finger (σ_2_-σ_4_ linker) that enters the RNAP active-center cleft and interacts with template-strand ssDNA (refs. 6-8, 44, 45). However, the σ fingers of ECF σ factors are only functional analogs--not also structural homologs--of the σ fingers of primary σ factors (refs. 1, 6). The σ fingers of ECF σ factors are shorter than, and are unrelated in sequence to, the σ fingers of primary σ factors (refs. 6-8). As a first step to assess whether the difference in σ-finger length and sequence for ECF σ factors vs. primary σ factors affects the mechanism of initiation-factor displacement, we have performed an analysis analogous to that in Fig. 1, but assessing ECF-σ-factor dependent, primer-dependent transcription initiation by *Mtb* RNAP holoenzyme containing the ECF σ factor σ^H^, which has a σ finger that reaches substantially less deeply--∼9 Å less deeply--into the RNAP active-center cleft than the σ finger of a primary σ factor (Fig. S4; refs. 6-8).

In previous work, one of our laboratories determined a crystal structure of a post-translocated-state initial transcribing complex comprising *Mtb* RNAP, the *Mtb* ECF σ factor σ^H^, promoter DNA, and a 5’-hydroxyl-containing RNA 6 nt in length (*Mtb σ*^H^-RPitc6; PDB 6JCX; ref. 6). Here, we determined crystal structures of post-translocated-state initial transcribing complexes containing 5’-hydroxyl-containing RNAs 5, 7, 9, and 10 nt in length (*Mtb σ*^H^-RPitc5, σ^H^-RPitc7, σ^H^-RPitc9, and σ^H^-RPitc10; Figs. 3A, C-E and S1C; Table S3). (We also attempted to determine a crystal structure of the corresponding complex containing a 5’-hydroxyl-containing RNA 8 nt in length, but this attempt was unsuccessful, instead yielding a structure of a pre-translocated-state complex.) Together with the published structure of *Mtb σ*^H^-RPitc6, the new structures provide a series of structures in which the position of the RNA 5’-end relative to RNAP varies, and in which the position of RNA 3’-end relative to RNAP remains constant, as in RNA extension in initial transcription (Fig. 3).

In the crystal structures of *Mtb σ*^H^-RPitc5, σ^H^-RPitc6, and σ^H^-RPitc7, the conformations of the σ^H^ finger are essentially identical (Figs. 3A-C). At least one of residues T106, T110, W112, and Q113 of the σ^H^ finger make H-bonded or van der Waals interactions with the template-strand DNA nucleotides at positions −6 and −7, pre-organizing the template strand to adopt a helical conformation and to engage the RNAP active center. In σ^H^-RPitc5 and σ^H^-RPitc6, there are no direct contacts between σ^H^ finger and the RNA. In σ^H^-RPitc7, there are direct contacts between the σ^H^ finger and the RNA 5’ end, with σ^H^ finger residues E107, Q108, and T110 contacting RNA nucleotide at positions −6 in a manner that likely stabilizes the RNA-DNA hybrid.

In contrast, in the crystal structures of *Mtb σ*^H^-RPitc9 and σ^H^-RPitc10, the σ^H^ finger adopts different conformations as compared to the structures of *Mtb σ*^H^-RPitc5, σ^H^-RPitc6, and σ^H^-RPitc7 (Figs. 3A-E), and the differences in conformation coincide with, and appear to be driven by, the movement of the 5’ end of RNA into the positions previously occupied by the tip of the σ finger and resulting steric collision between the RNA and the σ finger (Figs. 3A-E). In the structure of σ^H^-RPitc9, a segment comprising 20 residues of the σ^H^ finger--σ^H^ residues 102-121--exhibits segmental disorder, indicating that this 20-residue segment is flexible and exists as an ensemble of multiple conformations (Figs. 3D). In the structure of σ^H^-RPitc10, a segment comprising 32 residues of the σ^H^ finger--σ residues 94-125--exhibits segmental disorder, indicating that this 32-residue segment is flexible and exists as an ensemble of multiple conformations (Fig. 3E). Like the changes in conformation of the σ finger of the primary σ factor described in the preceding sections, the change in conformation of the σ^H^ finger can be understood as the folding of the loop of the σ-finger hairpin back over the strands of the σ-finger hairpin (Fig. 3A-E). We thus interpret the changes in conformation from σ^H^-RPitc7 to σ^H^-RPitc9 to σ^H^-RPitc10 as successive displacements of the loop of the σ-finger hairpin--involving displacement of first a 20-residue segment and then a 32-residue segment--back over the strands of the σ-finger hairpin, accompanied by increased flexibility of the displaced segments (Figs. 3A-D).

Taken together, the crystal structures of *Mtb σ*^H^-RPitc5, σ^H^-RPitc6, σ^H^-RPitc7, σ^H^-RPitc9, and σ^H^-RPitc10 suggest that nascent RNA with a 5’-hydroxyl end displaces the σ finger of ECF σ factor σ^H^ in a stepwise fashion, starting upon extension of nascent RNA to a length of 8 or 9 nt and continuing upon further extension through a length of at least 10 nt (Fig. 5C). We conclude that the mechanism of initiation-factor displacement for ECF σ factor σ^H^ is fundamentally similar to the mechanism of initiation-factor displacement for the primary σ factor, and we suggest that the later start of the process with ECF σ factor σ^H^ than with the primary σ factor--displacement starting at an RNA length of 8 or 9 nt, vs. displacement starting at an RNA length of 5 nt--is a consequence of the fact that the σ finger of σ^H^ finger reaches substantially less deeply--∼9 Å less deeply--into the RNAP active-center cleft than the σ finger of the primary σ factor (Fig. S4A-B)

**Figure 5.**
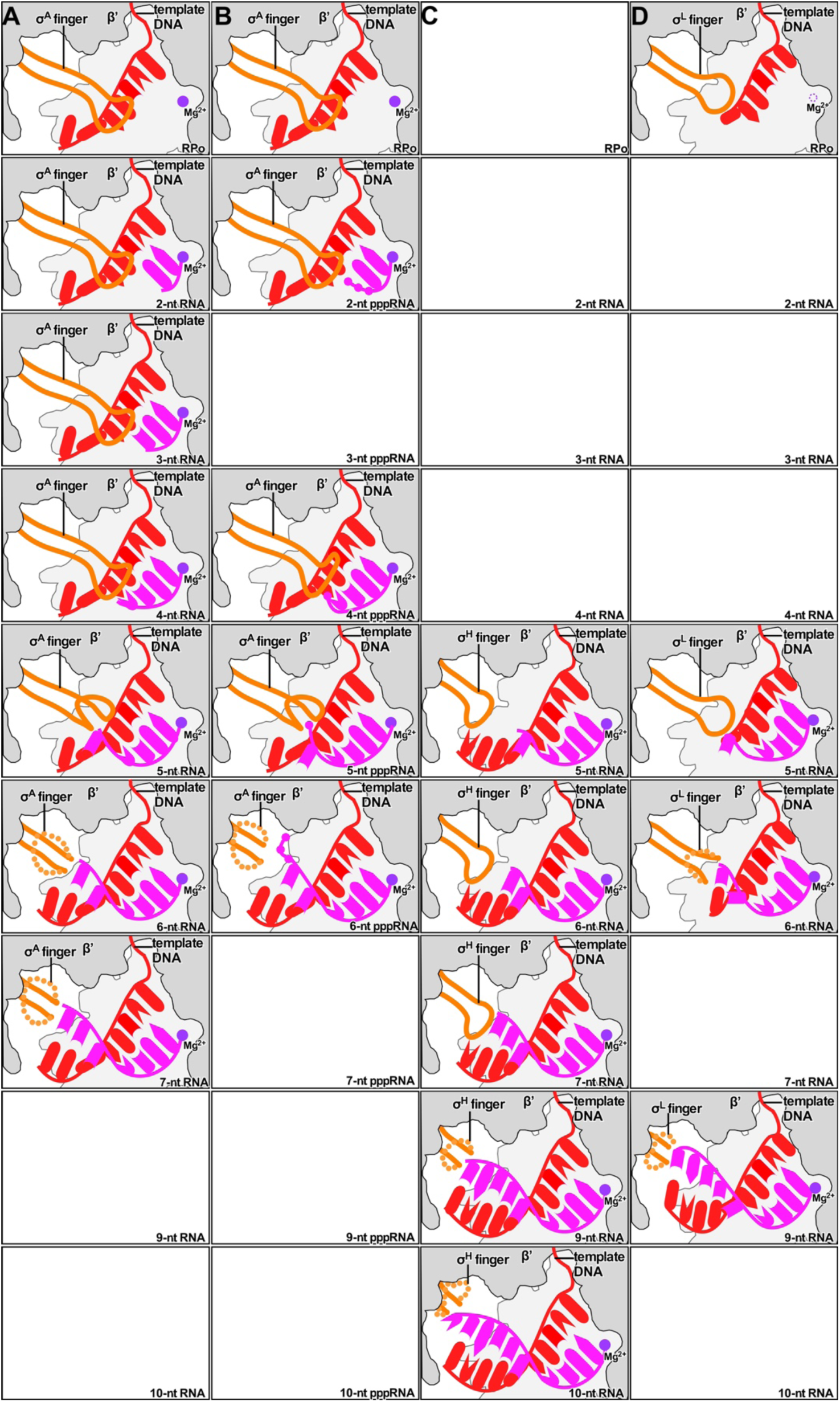
Summary. Schematic illustration of stepwise, RNA-driven displacement of σ finger with **(A)** primary σ factor in primer-dependent transcription initiation (Fig. 1), **(B)** primary σ factor in primer-independent transcription initiation (Fig. 2), **(C)** ECF σ factor σ^H^ (Fig. 3), and **(D)** ECF σ factor σ^L^ (Fig. 4). Colors as in Fig. 1. Disordered σ-finger segments are shown as dotted orange line, and disordered 5’-triphospahe-RNA phosphates are omitted.

### Stepwise, RNA-driven displacement of the σ finger: ECF σ factor σ^L^

The σ fingers of different ECF σ factors vary in length and sequence (ref. 6). As a second step to assess whether the difference in σ-finger length and sequence for ECF σ factors vs. primary σ factors affects the mechanism of initiation-factor displacement, we have performed an analysis analogous to that in Fig. 1, but assessing ECF-σ-factor dependent, primer-dependent transcription initiation by *Mtb* RNAP holoenzyme containing the ECF σ factor, σ^L^, which has a σ finger that reaches slightly less deeply--∼5 Å less deeply--into the RNAP active-center cleft than the σ finger of a primary σ factor (Fig. S4A-C; ref. 7).

In previous work, one of our laboratories determined crystal structures of a transcription initiation complex comprising *Mtb* RNAP, the *Mtb* ECF σ factor σ^L^, and promoter DNA (*Mtb σ*^L^-RPo; PDB 6DVE; ref. 7) and of a post-translocated-state initial transcribing complex comprising *Mtb* RNAP, the *Mtb* ECF σ factor σ^L^, promoter DNA, and a 5’-hydroxyl-containing RNA 5 nt in length (*Mtb σ*^L^-RPitc5; PDB 6DVB; ref. 7). Here, we determined a higher-resolution crystal structure of a post-translocated-state initial transcribing complex having a 5’-hydroxyl-containing RNA 5 nt in length (*Mtb σ*^L^-RPitc5) and crystal structures of post-translocated-state initial transcribing complexes containing 5’-hydroxyl-containing RNAs 6 and 9 nt in length (*Mtb σ*^L^-RPitc6 and σ^L^-RPitc9; Figs. 4B-D and S1D; Table S4). (We also attempted to determine crystal structures of the corresponding complex containing 5’-hydroxyl-containing RNAs 7 and 8 nt in length, but these attempts were unsuccessful, instead yielding structures of pre-translocated-state complexes.) Together with the published structures of *Mtb σ*^L^-RPo and σ^L^-RPitc5, the new structures provide a series of structures in which the position of the RNA 5’-end relative to RNAP varies, and in which the position of RNA 3’-end relative to RNAP remains constant, as in RNA extension in initial transcription (Fig. 4).

In the crystal structures of *Mtb σ*^L^-RPo and σ^L^-RPitc5, the conformations of the σ^L^ finger are identical (Figs. 4A-B; ref. 7). In the crystal structure of *Mtb σ*^L^-RPo, template-strand ssDNA was not resolved (Fig. 4A; ref. 7). In the crystal structure of *Mtb σ*^L^-RPitc5, residue S96 of the σ^L^ finger makes H-bonded interaction with the template-strand DNA nucleotide at position −5, pre-organizing the template strand to adopt a helical conformation and to engage the RNAP active center (Fig. 4B; ref. 7).

In contrast, in the crystal structures of *Mtb σ*^L^-RPitc6 and σ^L^-RPitc9, the σ^L^ finger adopts different conformations as compared to the structures of *Mtb σ*^L^-RPo and σ^L^-RPitc5 (Figs. 4A-D), and the differences in conformation coincide with, and appear to be driven by, the movement of the 5’ end of RNA into the positions previously occupied by the tip of the σ finger and resulting steric collision between the RNA and the σ finger (Figs. 4A-D). In the structure of σ^L^-RPitc6, two residues of the σ^L^ finger--σ^L^ residues 96-97--exhibit segmental disorder, indicating that this two-residue segment is flexible and exists as an ensemble of multiple conformations (Fig. 4C). In the structure of σ^L^-RPitc9, fifteen residues of the σ^L^ finger--σ residues 86-100--exhibit segmental disorder, indicating that this fifteen-residue segment is flexible and exists as an ensemble of multiple conformations (Fig. 4D). Like the change in conformation of the σ finger of the primary σ factor and the σ^H^ finger described in the preceding sections, the change in conformation of the σ^L^ finger can be understood as the folding of the loop of the σ-finger hairpin back over the strands of the σ-finger hairpin (Fig. 4B-D). We thus interpret the changes in conformation from σ^L^-RPitc5 to σ^L^-RPitc6 to σ^L^-RPitc9 as successive displacements of the loop of the σ-finger hairpin--involving displacement of first a two-residue segment and then a fifteen-residue segment--back over the strands of the σ-finger hairpin, accompanied by increased flexibility of the displaced segments (Figs. 4B-D).

Taken together, the crystal structures of *Mtb σ*^L^-RPo, σ^L^-RPitc5, σ^L^-RPitc6, and σ^L^-RPitc9 suggest that nascent RNA with a 5’-hydroxyl end displaces the σ finger of ECF σ factor σ^L^ in a stepwise fashion, starting upon extension of nascent RNA to a length of 6 nt and continuing upon further extension through a length of at least 9 nt. We conclude that the mechanism of initiation-factor displacement for ECF σ factor σ^L^ is fundamentally similar to the mechanism of initiation-factor displacement for the primary σ factor, and we suggest that the slight difference in starting points of the process with σ^L^, and the primary σ factor--displacement starting at an RNA length of 6 nt, vs. displacement starting at an RNA length of 5 nt--is a consequence of the fact that the σ finger of σ^L^ reaches slightly less deeply--∼5 Å less deeply--into the RNAP active-center cleft than the σ finger of the primary σ factor (Fig. S4).

### Mechanistic conclusions

For both the primary σ factor and the studied ECF σ factors, the portion of the σ finger that interacts with template-strand ssDNA in the RNAP-promoter open complex and is displaced in the RNAP-promoter initial transcribing complex is an unstructured segment that contains, at its tip, a loop (Figs. 1-4; summary in Fig. 5). For σ^A^, with both 5’-hydroxyl and 5’-triphosphate nascent RNAs, the first step in displacement of the σ finger is folding of the loop back on itself (Figs. 1D-E, 2C-D, and Fig. 5A-B), and successive steps in displacement, which are associated with successive increases in disorder of the loop, are best interpreted as successive steps of folding of the loop further back on itself (1D-G, 2C-E, and Fig. 5A-B). For ECF σ factors, the first and successive steps in displacement of the σ finger entail successive disorder of the loop at the tip of the σ finger and likewise are best interpreted as successive folding of the loop back on itself (Figs. 3C-E, 4B-D, and 5C-D). We point out that a protein segment that forms a hairpin loop is particular well suited for stepwise folding back on itself and thus is particular well suited for stepwise storage of “stress energy” (Fig. 5). We suggest that such a segment may serve as a “protein spring” that is compressed in stepwise fashion during initial transcription, and that this “protein spring” may contribute--together with a “DNA spring” entailing DNA scrunching (refs. 2, 46, 47)--towards stepwise storage of energy required for subsequent breakage of RNAP-promoter and RNAP-σ interactions in promoter escape.

Our results show that the σ finger plays different roles during different stages of initial transcription (Fig. 5). In early stages of initial transcription, the σ finger functions as an RNA mimic, pre-organizing template-strand ssDNA, first to engage the RNAP active center to enable binding of the initiating and extending NTPs, and then to stabilize the short RNA-DNA hybrids formed by 2-4 nt RNA products. In contrast, in later stages of initial transcription, the σ finger poses a steric and energetic barrier for RNA extension, and collision of the RNA 5’ end with the σ finger disrupts interactions of the σ finger with template-strand ssDNA, drives stepwise displacement of the σ finger, and potentially drives stepwise compression of a σ-finger “protein spring,” thereby potentially storing energy for use in subsequent breakage of RNAP-promoter and RNAP-σ interactions in promoter escape.

Although we have analyzed only the primary σ factor and two ECF alternative σ factors, we note that other alternative σ factors also have a structural module that enters the active-center cleft, and interacts with template ssDNA in manner that blocks the path of nascent RNA (refs. 1, 14). Non-ECF alternative σ factors other than σ^54^ typically have a σ finger closely similar in sequence to the σ finger of the primary σ factor (refs. 1, 14). We predict that non-ECF alternative σ factors other than σ^54^ will exhibit the same, or a closely similar, mechanism of stepwise, RNA-driven displacement as for the primary σ factor, with displacement starting at the same, or a closely similar, RNA length as for the primary σ factor (i.e., ∼5 nt RNA in post-translocated state; ∼6 nt RNA in pre-translocated state). σ^54^ is structurally unrelated to the primary σ factor and other alternative σ factors, but, nevertheless, has a structural module, termed RII.3, that contains a loop that makes analogous interactions with template-strand ssDNA (refs. 13, 15). We predict that σ^54^ will exhibit an analogous mechanism of stepwise, RNA-driven displacement, and, based on structural modelling, we predict that displacement will start at the same, or a similar, RNA length as for the primary σ factor (i.e., ∼5 nt RNA in post-translocated state; ∼6 nt RNA in pre-translocated state; Fig. S5A-C).

Archaeal-RNAP and eukaryotic RNAP-I, RNAP-II, and RNAP-III transcription initiation complexes also contain structural modules--homologous to each other, but not homologous to the relevant structural modules of bacterial σ factors--that enter the active-center cleft, and interact with template ssDNA in manner that blocks the path of nascent RNA (TFB zinc ribbon and CSB for archaeal RNAP; Rrn7 zinc ribbon and B-reader for RNAP I; TFIIB zinc ribbon and B-reader for RNAP II; and Brf1 zinc ribbon for RNAP III (refs. 16-18, 21-27, 30). We predict that these structural modules will exhibit an analogous mechanism of stepwise, RNA-driven displacement, and based on structural modelling, we predict that that displacement will start at a similar, RNA length as for the primary σ factor (i.e., ∼7 nt RNA in post-translocated state; ∼8 nt RNA in pre-translocated state; Fig. S6A-C).

## METHODS

### Structure determination

The assembly and crystallization of the complexes of Figs. 1-2 were performed by Y.Z. at Rutgers University, using procedures as in (ref. *5*). Crystals were obtained using a reservoir solution of 0.1 M Tris-HCl, pH 8.0, 200 mM KCl, 50 mM MgCl_2_, and 10% (m/v) PEG 4000. Structures were determined by the molecular replacement method.

The assembly and crystallization of the complexes of Fig. 3 were performed by L.L. at Shanghai Institute of Plant Physiology and Ecology, using procedures as in (ref. *6*). Crystals were obtained using a reservoir solution of 50 mM sodium cacodylate, pH 6.5, 80 mM Mg(OAc)_2_, and 15% PEG-400. Structures were determined by the molecular replacement method.

The assembly and crystallization of the complexes of Fig. 4 were performed by V.M. and W.L. at Rutgers University. Crystals were obtained in a reservoir solution of 100 mM sodium citrate, pH 5.6, 200 mM sodium acetate, and 10% (m/v) PEG 4000. Structures were determined by the molecular replacement method.

Detailed methods are provided in the Supporting Information file.

## DATA AVAILABILITY

Coordinates have been deposited in the PDB under ID codes 6KQD, 6KQE, 6KQF, 6KQG, and 6KQH for *Tt σ*^A^-RPitc3, *Tt σ*^A^-RPitc4, *Tt σ*^A^-RPitc5, *Tt σ*^A^-RPitc6, and *Tt σ*^A^-RPitc7, respectively; coordinates have been deposited in the PDB under ID codes 6L74, 6KQL, 6KQM, and 6KQN for *Tt σ*^A^-RPitc2-PPP, *Tt σ*^A^-RPitc4-PPP, *Tt σ*^A^-RPitc5-PPP, *Tt σ*^A^-RPitc6-PPP, respectively; coordinates have been deposited in the PDB under ID codes 6KON, 6KOO, 6KOP, and 6KOQ for *Mtb σ*^H^-RPitc5, *Mtb σ*^H^-RPitc7, *Mtb σ*^H^-RPitc9, and *Mtb σ*^H^-RPitc10, respectively; and coordinates have been deposited in the PDB under ID codes 6TYE, 6TYF, and 6TYG for *Mtb σ*^L^-RPitc5, *Mtb σ*^L^-RPitc6, and *Mtb σ*^L^-RPitc9, respectively.

## ACKNOWLEDGMENTS

This work was supported by Chinese Natural Science Foundation of China grant 31670067 and 31822001 to Y.Z., the CAS Leading Science Key Research Program grant QYZDB-SSW-SMC005 to Y.Z., and NIH grant GM041376 to R.H.E. We thank the Argonne National Laboratory Advanced Photon Source, the Brookhaven National Synchrotron Light Source II, the Cornell High Energy Synchrotron Source, and the Shanghai Synchrotron Radiation Facility for assistance during data collection.

## AUTHOR CONTRIBUTIONS

L.L., V.M., W.L., and Y.Z. determined structures. Y.Z. and R.H.E. designed experiments, analyzed data, and wrote the manuscript.

