## Supplemental text, methods, and figures for "RNA extension drives a stepwise displacement of an initiation-factor structural module in initial transcription"

**SUPPORTING INFOMATION**

***Thermus thermophilus* (*Tth*) $\sigma^A$**

The wild-type *Tth* $\sigma^A$ was prepared from *Escherichia coli* BL21(DE3) (Invitrogen, Inc.) transformed with plasmid pET28a-*Tth*$\sigma^A$, as in (ref. 1).

***Mycobacterium tuberculosis* (*Mtb*) RNAP $\sigma^H$**

*Mtb* $\sigma^H$ was prepared from *E. coli* BL21(DE3) (Novoprotein, Inc.) transformed with plasmid pTolo-EX5-*Mtb*$\sigma^H$ (ref. 2), as in (ref. 2).

***Tth* RNAP core enzyme**

*Tth* RNAP core enzyme was prepared from *Tth* strain HB8 (DSM579; Deutsche Sammlung von Mikroorganismen und Zellkulturen GmbH) as in (ref.1).

***Mtb* RNAP core enzyme**

*Mtb* RNAP core enzyme was prepared from *E. coli* BL21(DE3) (Novoprotein, Inc.) transformed with plasmids pETDuet-*Mtb*-rpoA-rpoZ (ref. 2) and pACYCDuet-*Mtb*-rpoB-rpoC (ref. 2), as in (ref. 2).

***Tth* RNAP $\sigma^A$ holoenzyme**

*Tth* RNAP holoenzyme was prepared as in (ref. 1). Briefly, *Tth* RNAP core enzyme (13 μM) and *Tth* $\sigma^A$ (52 μM) were incubated in 2 ml 20 mM Tris-HCl, pH 7.7, 150 mM NaCl, and 2% glycerol for 1 h at 4 °C. The mixture was applied to a HiLoad 16/60 Superdex S200 column (GE Healthcare, Inc.) equilibrated in a running buffer of 20 mM Tris-HCl, pH 7.7, 100 mM NaCl, and 1% glycerol, and the *Tth* RNAP holoenzyme was eluted with the same buffer. Fractions containing *Tth* RNAP holoenzyme were pooled, concentrated to ~7.5 mg/ml using 30 kDa MWCO Amicon Ultra-15 centrifugal ultrafilters (Millipore, Inc.), and stored in the same buffer at -80 °C.

### *Mtb* RNAP $\sigma^L$ holoenzyme

*Mtb* RNAP $\sigma^L$ holoenzyme was prepared from *E. coli* strain BL21(DE3)STAR (Invitrogen, Inc.) transformed with plasmids pCOLADuet-*rpoB*-*rpoC (3-4)*, pACYC-*rpoA*-*sigL* [encodes *Mtb* RNAP $\alpha$ subunit and octahistidine-tagged *Mtb* $\sigma^L$; constructed from plamid pACYCDuet-*rpoA*(ref. 4) by deletion of the descahistidine-tag coding seqence preceding *rpoA* by use of site-directed mutagenesis (QuikChange II Site-Directed Mutagesesis Kit; Agilent, Inc.) and replacment of the BglII-XhoI DNA segment by the BglII-XhoI DNA segment of a DNA fragment, comprising 5'-ATGGCAGATCTAATGCATCATCATCATCATCATCATCAT-3' followed by codons 2-177 of *Mtb* $\sigma^L$ followed by 5'- TGACTCGAG-3', generated by PCR using plasmid pSR32 (ref. 5) as template], and pCDF-*rpoZ* (ref. 4). Single colonies of the resulting transformants were used to inoculate 50 ml LB broth containing 100 μg/ml ampicillin, 35 μg/ml chloramphenicol, and 100 μg/ml streptomycin; cultures were incubated at 37°C with shaking until $OD_{600}$ = 0.7; cultures were induced by addition of IPTG to 1 mM , and cultures were further incubated at 16 h 18°C with shaking. Cells were harvested by centrifugation (4,000xg; 10 min at 4°C), re-suspended in 50 ml buffer A (20 mM Tris-HCl, pH 8.0, 200 mM NaCl, 2 mM $ZnCl_2$, 1 mM EDTA, 5 mM 2-mercaptoethanol, and 5% glycerol) containing 1 tablet cOmplete Protease Inhibitor Cocktail (Roche, Inc.) at 4°C on ice; and lysed by sonication [4 min on ice; 4 s pulses at 40 W with 2 s intervals between pulses using Branson Ultrasonics Sonifier S-450A (Branson Ultrasonics, Inc.)]. The lysate was cleared by centrifugation (26,000 x g; 30 min at 4°C), and the cleared lysate (40 ml) was mixed with 2.8 ml 0.7% polyethyleneimine (Polymin-P; Sigma-Aldrich, Inc.), incubated 1 h with rotary shaking, and centrifuged (26,000xg; 10 min 4ºC). The polyethylene precipitate was washed 3 times by re-suspension in 40 ml buffer B (20 mM Tris-HCl, pH 8.0, 0.1 mM EDTA, 5 mM 2-mercaptoethanol, and 5% glycerol) containing 500 mM NaCl at 4°C, followed by centrifugation (26,000xg; 10 min at 4ºC), and was extracted with 20 ml buffer B containing 1 M NaCl at 4°C. The extract was cleared by centrifugation (26,000xg; 10 min at 4ºC), and the cleared extract was mixed with 30 ml ice-cold saturated ammonium sulfate, incubated 1 h with rotary shaking, and centrifuged (26,000xg; 30 min at 4ºC). The ammonium sulfate precipitate was dissolved in 30 ml buffer B containing 200 mM NaCl at 4°C, dialysed 16 h at

4°C against 2 L buffer B containing 200 mM NaCl, and an aliquot (30 ml) was applied to a 10 ml column of Ni-NTA agarose column (Qiagen, Inc.; pre-equilibrated in buffer B containing 200 mM NaCl; 4°C). The column was washed with 150 ml buffer B containing 200 mM NaCl and 15 mM imidazole, and was eluted with 20 ml buffer B containing 200 mM NaCl and 200 mM imidazole. The eluate was applied to two tandem 5 ml HiTrap Q HP columns (GE Healthcare, Inc.; pre-equilibrated with buffer B containing 200 mM NaCl; 4°C; flow rate = 0.2 ml/min), the column was washed with 100 ml buffer B containing 200 mM NaCl, and the column was eluted with 200 ml of a linear gradient of 200-600 mM NaCl in buffer B at 4°C, and pooled fractions containing Mtb RNAP $\sigma^L$ holoenzyme were concentrated to 5 ml using 30 kDa MWCO Amicon Ultra-15 centrifugal ultrafilters (Millipore, Inc.). The sample was further purified by gel filtration on a HiLoad 16/60 Superdex 200 prep grade column (GE Healthcare, Inc) in 20 mM Tris-HCl, pH 8.0, 75 mM NaCl, 5 mM $MgCl_2$, and 5 mM dithiothreitol, concentrated to 20 μM in the same buffer using 30 kDa MWCO Amicon Ultra-15 centrifugal ultrafilters, flash-frozen in liquid N2, and stored at -80°C. Yields were ~5 mg/L, and purities were >95%.

**Nucleic-acid scaffolds**

Oligodeoxyribonucleotides (IDT for *Tth* $\sigma^A$-RPitcs, *Tth* $\sigma^A$-RPitcs-PPP, and *Mtb* $\sigma^L$-RPitc; Sangon Biotech for *Mtb* $\sigma^H$-RPitcs) and oligoribonucleotides (IDT for structures of *Tth* $\sigma^A$-RPitc, *Tth* $\sigma^A$-RPitc-PPP, and *Mtb* $\sigma^L$-RPitc; GenScript Biotech for *Mtb* $\sigma^H$-RPitc) were dissolved in ultrapure water (GIBCO) to 3 mM and stored at -80°C.

Nucleic-acid scaffolds (sequences in Fig. S1) for structures in Fig. 1-3 were prepared as follows. Nontemplate-strand DNA (0.5 mM final), template-strand DNA (0.55 mM final), and RNA (1 mM final; omitted for *Tth* $\sigma^A$-RPitc2-PPP) were combined in 25 μl 5 mM Tris-HCl, pH 7.7, 200 mM NaCl, and 10 mM $MgCl_2$; were heated 5 min at 95°C; were cooled to 25°C in 2°C steps with 1 min/step using a thermal cycler (Applied Biosystems); and were stored at -80 °C.

Nucleic-acid scaffolds (sequences in Fig. S1) for structures in Fig. 4 were prepared as follows. Nontemplate-strand DNA (1 mM final), template-strand DNA (1 mM final), and RNA (1 mM final) were combined in 50 μl 5 mM Tris-HCl, pH 7.5; were heated 8 min at

95°C;  cool to 22°C over 2 h, and were stored at -80°C.

**Assembly of transcription initiation complexes**

For structures in Figs. 1-2, complexes for crystallization were prepared by mixing 20 μl 18 μM *Tth* $\sigma^A$ holoenzyme (in 20 mM Tris-HCl, pH 7.7, 100 mM NaCl, and 1% glycerol), 1 μl 0.5 mM nucleic-acid scaffold (in 5 mM Tris-HCl, pH 7.7, 200 mM NaCl, and 10 mM $MgCl_2$), and 1 μl 25 nM 5'-triphosphate-GpA (in water; *Tth* $\sigma^A$-RPitc2-PPP only), and incubating 1 h at 25°C.

For structures in Fig. 3, complexes for crystallization were prepared by mixing 500 μl 20 μM *Mtb* RNAP core enzyme (in 20 mM Tris-HCl, pH 7.7, 100 mM NaCl, and 1% glycerol), 120 μl 330 μM *Mtb* $\sigma^H$, and 30 μl 0.5 mM nucleic-acid scaffold (in 5 mM Tris-HCl, pH 7.7, 200 mM NaCl, and 10 mM $MgCl_2$), and incubating 12 h at 4 °C. The sample was loaded onto a Hiload 16/60 Superdex S200 column (GE Healthcare, Inc.) equilibrated in 10 mM Tris-HCl pH 8.0, 0.1 M NaCl, 1% (v/v) glycerol, 1 mM dithiothreitol at 4°C and eluted with 120 ml of the same buffer at 4°C. Pooled fractions containing *Mtb* $\sigma^H$-RPitc were concentrated to 7.5 mg/ml using 100 kDa MWCO Amicon Ultra-15 centrifugal ultrafilters (Millipore, Inc.) at 4°C.

For structures in Fig. 4, complexes for crystallization were prepared by mixing 20 μl 20 μM *Mtb* RNAP $\sigma^L$ holoenzyme (in 20 mM Tris-HCl, pH 8.0, 75 mM NaCl, 5 mM $MgCl_2$, and 5 mM dithiothreitol) with 2 μl 1 mM nucleic-acid scaffold (in 5 mM Tris-HCl, pH 7.5), and incubating 20 min at 22°C.

**Crystallization, data collection, and structure determination: *Tth* $\sigma^A$-RPitc and $\sigma^A$-RPitc-PPP**

For structures in Figs. 1-2, crystallization was performed as in (ref. 1), but using the complexes of the preceding section. Briefly, crystallization drops containing 1 μl $\sigma^A$-RPitc (20 μM in 10 mM Tris-HCl, pH8.0, 100 mM NaCl, 0.5% glycerol, 1mM DTT), and 1 μl reservoir solution A [0.1 M Tris-HCl, pH8.0, 200 mM KCl, 50 mM $MgCl_2$, and 10% (m/v) PEG 4000] were equilibrated to 400 μl reservoir solution A in sealed hanging-drop plates.

Crystals were grown 3 days at 22°C. Micro-seeding was performed as needed using the same reservoir solution. Crystals were transferred to reservoir solution A containing 18% (v/v) (2R, 3R)-(-)-2,3-butanediol (Sigma Aldrich) and flash-cooled in liquid nitrogen.

Diffraction data were collected at synchrotron beamlines CHESS-F1 and BNL-X25 and were processed using HKL2000 (ref. 6). Structures were solved by molecular replacement using the crystal structure of *Tth* RPo (PDB 4G7H; ref. 1) as search model. Iterative cycles of reciprocal refinement using Phenix (ref. 7) and real-space model building using Coot (ref. 8) were performed. The final atomic models and structure factors were deposited in the Protein Data Bank (PDB) with accession codes 6KQD, 6KQE, 6KQF, 6KQG, and 6KQH for *Tth* $\sigma^A$-RPitc3, $\sigma^A$-RPitc4, $\sigma^A$-RPitc5, $\sigma^A$-RPitc6, and $\sigma^A$-RPitc7, and with accession codes 6L74, 6KQL, 6KQM, and 6KQN for *Tth* $\sigma^A$-RPitc2-PPP, $\sigma^A$-RPitc4-PPP, $\sigma^A$-RPitc5-PPP, and $\sigma^A$-RPitc6-PPP, respectively (Tables S1-S2).

**Crystallization, data collection, and structure determination: *Mtb* $\sigma^H$-RPitc**

For structures in Fig. 3, crystallization was performed by mixing 1 μl *Mtb* $\sigma^H$-RPitc (20 μM in 10 mM Tris-HCl, pH8.0, 100 mM NaCl, 1% (v/v) glycerol, 1mM DTT) with 1 μl reservoir solution B [50 mM sodium cacodylate, pH 6.5, 80 mM magnesium cetate, and 15% (m/v) PEG 400]. Crystals were grown 1 week at 22°C in sealed hanging-drop plates, were cryo-protected in reservoir solution B containing 30% PEG 400, and were flash-cooled in liquid nitrogen.

Diffraction data were collected at Shanghai Synchrotron Radiation Facility (SSRF) beamlines 17U or 19U, and were processed using HKL2000 *(6)*. Structures were solved by molecular replacement using the crystal structure of *Mtb* $\sigma^H$-RPo (PDB 5ZX2; ref. 2) as search model. Iterative cycles of reciprocal refinement using Phenix (ref. 7) and real-space model building using Coot (ref. 8) were performed. The final atomic models and structure factors were deposited in the PDB with accession codes 6KON, 6KOO, 6KOP, and 6KOQ for *Mtb* $\sigma^H$-RPitc5, $\sigma^H$-RPitc7, $\sigma^H$-RPitc9, and $\sigma^H$-RPitc10, respectively (Table S3).

**Crystallization, data collection, and structure determination: *Mtb* $\sigma^L$-RPitc**

For structures in Fig. 4, crystallization of *Mtb* $\sigma^L$-RPitc was performed by mixing 1 µl *Mtb* $\sigma^L$-RPitc (20 µM in 20 mM Tris-HCl, pH 8.0, 75 mM NaCl, 5 mM $MgCl_2$, and 5 mM dithiothreitol) with 1 µl reservoir solution C [100 mM sodium citrate, pH 5.6, 200 mM sodium acetate, and (m/v) 10% PEG 4000]. Crystals were grown at 1 week at 22°C in sealed hanging-drop plates, were cryo-protected in reservoir solution containing 20% (v/v) (2R, 3R)-(-)-2,3-butanediol, and were flash-cooled in liquid nitrogen.

Diffraction data were collected at Argonne Photon Source (APS) beamline 19-BM. Structures were solved by molecular replacement usng the crystal structure of *Mtb* $\sigma^L$-$RP_O$ (PDB 6DVC; *8*) as search model. One molecule of RNAP was present per asymmetric unit. Cycles of nodel building and refinement were performed using Coot (ref. 8) and Phenix (ref. 7). The final models were generated by X/Y/Z-coordinate refinement using secondary-structure restraints, followed by B-factor refinement and individual-B-factor refinement. The final atomic models and structure factors were deposited in the PDB with accession codes 6TYE, 6TYF, and 6TYG for *Mtb* $\sigma^L$-RPitc5, $\sigma^L$-RPitc6, and $\sigma^L$-RPitc9, respectively (Table S4).

### SUPPLEMENTAL TABLES

Table S1. Structure-determination and refinement statistics: *Tth* $\sigma^A$-RPitc

| | ***Tt*** **$\sigma^A$-RPitc3** | ***Tt*** **$\sigma^A$-RPitc4** | ***Tt*** **$\sigma^A$-RPitc5** | ***Tt*** **$\sigma^A$-RPitc6** | ***Tt*** **$\sigma^A$-RPitc7** |
|---|---|---|---|---|---|
| **Data collection** | | | | | |
| Space group | $P2_1$ | C2 | C2 | C2 | C2 |
| Cell dimensions | | | | | |
| a, b, c (Å) | 185.5, 104.0, 297.4 | 184.6, 103.8, 296.0 | 183.6, 103.6, 296.2 | 184.2, 104.0, 296.8 | 183.0, 103.6, 294.9 |
| α, β, γ (°) | 90.0, 98.5, 90.0 | 90.0, 98.7, 90.0 | 90.0, 98.9, 90.0 | 90.0, 98.6, 90.0 | 90.0, 99.2, 90.0 |
| Resolution (Å) | 50.00-3.30 | 50.00-3.30 | 50.00-2.45 | 50.0-2.80 | 40.0-3.20 |
| $R_{sym}$ or $R_{merge}$ | 9.2 (51.0) | 13.2 (95.5) | 9.2 (82.6) | 7.4 (71.0) | 18.5 (86.3) |
| I/σI | 9.9 (1.6) | 9.5 (1.2) | 14.3 (1.7) | 16.3 (1.4) | 6.5/1.4 |
| Completeness | 0.916 (0.936) | 1.00 (0.997) | 0.999(0.998) | 0.998 (0.980) | 0.937 (0.911) |
| Redundancy | 2.8 (2.7) | 3.4 (3.3) | 4.3 (3.5) | 3.4 (3.0) | 3.5 (3.3) |
| **Refinement** | | | | | |
| Resolution (Å) | 50.00-3.30 | 50.00-3.30 | 50.00-2.45 | 50.00-2.80 | 40.00-3.20 |
| No. reflections | 154582 | 83410 | 201239 | 136907 | 85533 |
| Rwork/ Rfree | 20.7/24.9 | 21.8/25.8 | 19.9 (23.0) | 20.5 (24.8) | 20.3 (24.7) |
| No. of atoms | 57053 | 28598 | 28634 | 28596 | 28589 |
| Protein | 55239 | 27671 | 27687 | 27607 | 27580 |
| Nucleic acids | 1825 | 932 | 953 | 995 | 1015 |
| B-factors (Å$^2$) | 83.1 | 96.0 | 64.6 | 88.6 | 64.7 |
| Protein | 82.3 | 95.2 | 63.6 | 87.9 | 63.8 |
| Nucleic acids | 107.7 | 120.9 | 94.4 | 108.7 | 87.8 |
| R.m.s deviations | | | | | |
| Bond lengths (Å) | 0.003 | 0.008 | 0.015 | 0.005 | 0.008 |
| Bond angles (°) | 0.617 | 1.074 | 1.026 | 0.735 | 0.632 |
| Ramachandran plot | | | | | |
| Favored (%) | 98.5 | 97.5 | 97.6 | 98.6 | 98.8 |
| Allowed (%) | 1.5 | 2.5 | 2.4 | 1.4 | 1.2 |
| Disallowed (%) | 0 | 0 | 0 | 0 | 0 |
| PDB code | 6KQD | 6KQE | 6KQF | 6KQG | 6KQH |

Numbers in parentheses are for highest resolution.

Table S2. Structure-determination and refinement statistics: *Tth* $\sigma^A$-RPitc-PPP

| | ***Tt*** **$\sigma^A$-RPitc2-PPP** | ***Tt*** **$\sigma^A$-RPitc4-PPP** | ***Tt*** **$\sigma^A$-RPitc5-PPP** | ***Tt*** **$\sigma^A$-RPitc6-PPP** |
|---|---|---|---|---|
| **Data collection** | | | | |
| Space group | C2 | C2 | C2 | *C2* |
| Cell dimensions | | | | |
| a, b, c (Å) | 183.1, 103.5, 295.1 | 183.8, 103.2, 295.6 | 184.7, 102.7, 295.7 | 184.6, 101.9, 296.1 |
| α, β, γ (°) | 90.0, 99.2, 90.0 | 90.0, 99.1, 90.0 | 90.0, 98.9, 90.0 | 90.0, 98.8, 90.0 |
| Resolution (Å) | 45.00-3.10 | 45.00-2.88 | 50.00-3.20 | 50.00-3.50 |
| $R_{sym}$ or $R_{merge}$ | 14.1 (74.4) | 11.0 (48.9) | 14.1 (98.2) | 13.7 (71.5) |
| I/σI | 9.6 (1.6) | 11.3 (2.4) | 11.3 (1.5) | 9.7 (1.5) |
| Completeness | 0.999 (0.99) | 0.999 (0.991) | 0.998 (0.976) | 0.950 (0.789) |
| Redundancy | 3.6 (3.3) | 3.7 (3.5) | 5.2 (4.3) | 4.6 (3.6) |
| **Refinement** | | | | |
| Resolution (Å) | 45.00-3.10 | 45.00-2.90 | 50.00-3.20 | 50.00-3.50 |
| No. reflections | 96614 | 121225 | 86387 | 64854 |
| Rwork/ Rfree | 19.1/23.6 | 20.7/24.6 | 20.5/25.2 | 21.3/25.8 |
| No. of atoms | 28744 | 28609 | 28642 | 28612 |
| Protein | 27685 | 27678 | 27691 | 27611 |
| Nucleic acids | 877 | 937 | 957 | 1007 |
| B-factors ($Å^2$) | 89.1 | 82.2 | 50.9 | 128.3 |
| Protein | 88.2 | 81.4 | 50.7 | 127.9 |
| Nucleic acids | 122.5 | 106.4 | 58.6 | 138.8 |
| R.m.s deviations | | | | |
| Bond lengths (Å) | 0.009 | 0.010 | 0.008 | 0.008 |
| Bond angles (°) | 0.583 | 1.040 | 1.065 | 0.998 |
| Ramachandran plot | | | | |
| Favored (%) | 98.1 | 98.0 | 97.4 | 97.7 |
| Allowed (%) | 1.9 | 2.0 | 2.6 | 2.3 |
| Disallowed (%) | 0 | 0 | 0 | 0 |
| PDB code | 6L74 | 6KQL | 6KQM | 6KQN |

Numbers in parentheses are for highest resolution.

Table S3. Structure-determination and refinement statistics: *Mtb* $\sigma^H$-RPitc

| | ***Mtb*** $\sigma^H$**-RPitc5** | ***Mtb*** $\sigma^H$**-RPitc7** | ***Mtb*** $\sigma^H$**-RPitc9** | ***Mtb*** $\sigma^H$**-RPitc10** |
|---|---|---|---|---|
| **Data collection** | | | | |
| Space group | P21 | P21 | P21 | P21 |
| Cell dimensions | | | | |
| a, b, c (Å) | 129.3, 161.2, 129.5 | 125.0, 162.3, 128.5 | 127.4, 162.6, 133.4 | 126.9, 161.1, 129.6 |
| α, β, γ (°) | 90.0, 117.8, 90.0 | 90.0, 117.0, 90.0 | 90.0, 117.9, 90.0 | 90.0, 117.4, 90.0 |
| Resolution (Å) | 50.00-3.00 | 50.00-2.80 | 50.00-3.30 | 50-3.35 |
| Rsym or Rmerge | 10.5(107.2) | 11.1(102.0) | 11.9(154.7) | 14.4(97.9) |
| I/σI | 12.0(1.11) | 14.4(1.1) | 13.4(1.1) | 10.9(1.2) |
| Completeness | 0.978(0.88) | 0.975(0.831) | 0.986(0.992) | 0.977(0.850) |
| Redundancy | 4.2(3.6) | 5.5(4.7) | 4.2(4.2) | 5.4(4.2) |
| CC1/2 in highest shell | 0.506 | 0.608 | 0.502 | 0.600 |
| **Refinement** | | | | |
| Resolution (Å) | 50.00-3.00 | 50.00-2.80 | 50.00-3.30 | 50.00-3.35 |
| No. reflections | 91412 | 109363 | 70820 | 64242 |
| Rwork/ Rfree | 22.1/27.1 | 20.6/24.8 | 21.7/25.5 | 22.1/26.0 |
| No. of atoms | 24039 | 24837 | 24560 | 24473 |
| Protein | 23117 | 23779 | 23450 | 23370 |
| Nucleic acids | 897 | 1029 | 1093 | 1082 |
| B-factors (Å2) | 95.42 | 88.28 | 121.84 | 103.06 |
| Protein | 94.08 | 87.80 | 121.78 | 102.44 |
| Nucleic acids | 130.47 | 99.94 | 125.3.71 | 117.02 |
| R.m.s deviations | | | | |
| Bond lengths (Å) | 0.003 | 0.006 | 0.002 | 0.004 |
| Bond angles (°) | 0.554 | 0.765 | 0.514 | 0.661 |
| Ramachandran plot | | | | |
| Favored (%) | 94.6 | 96.64 | 97.0 | 97.0 |
| Allowed (%) | 5.4 | 3.6 | 3.0 | 3.0 |
| Disallowed (%) | 0 | 0 | 0 | 0 |
| PDB code | 6KON | 6KOO | 6KOP | 6KOQ |

Numbers in parentheses are for highest resolution.

Table S4. Structure-determination and refinement statistics: *Mtb* $\sigma^L$-RPitcs

| | ***Mtb* $\sigma^L$-RPitc5** | ***Mtb* $\sigma^L$-RPitc6** | ***Mtb* $\sigma^L$-RPitc9** |
|---|---|---|---|
| **Data collection** | | | |
| Space group | P212121 | P212121 | P212121 |
| Cell dimensions | | | |
| a, b, c (Å) | 129.5, 158.5, 214.6 | 126.4, 161.7, 213.9 | 128.6, 158.2, 215.2 |
| α, β, γ (°) | 90.0, 90.0, 90.0 | 90.0, 90.0, 90.0 | 90.0, 90.0, 90.0 |
| Resolution (Å) | 48.01-3.30 | 49.26-3.30 | 49.12-2.92 |
| Rsym or Rmerge | 25.3(102.4) | 18.6(89.4) | 11.4(28.8) |
| I/σI | 9.1(2.4) | 5.6(1.1) | 7.4(6.7) |
| Completeness | 0.99(1.0) | 0.95(0.98) | 0.99(0.99) |
| Redundancy | 9.0(8.7) | 8.3(8.5) | 6.8(8.6) |
| CC1/2 in highest shell | 0.879 | 0.622 | 0.976 |
| **Refinement** | | | |
| Resolution (Å) | 48.01-3.80 | 49.26-3.80 | 49.12-3.50 |
| No. reflections | 42598 | 42516 | 55264 |
| Rwork/ Rfree | 25.2/30.3 | 24.8/30.6 | 24.1/29.5 |
| No. of atoms | 24794 | 24825 | 24818 |
| Protein | 23865 | 23834 | 23745 |
| Nucleic acids | 929 | 991 | 1073 |
| B-factors (Å2) | 91.66 | 89.54 | 73.09 |
| R.m.s deviations | | | |
| Bond lengths (Å) | 0.003 | 0.006 | 0.014 |
| Bond angles (°) | 0.635 | 0.768 | 1.149 |
| Ramachandran plot | | | |
| Favored (%) | 93.83 | 95.16 | 93.99 |
| Allowed (%) | 5.16 | 4.39 | 5.06 |
| Disallowed (%) | 1.01 | 0.46 | 0.95 |
| PDB code | 6TYE | 6TYF | 6TYG |

Numbers in parentheses are for highest resolution.

### SUPPLEMENTAL FIGURES

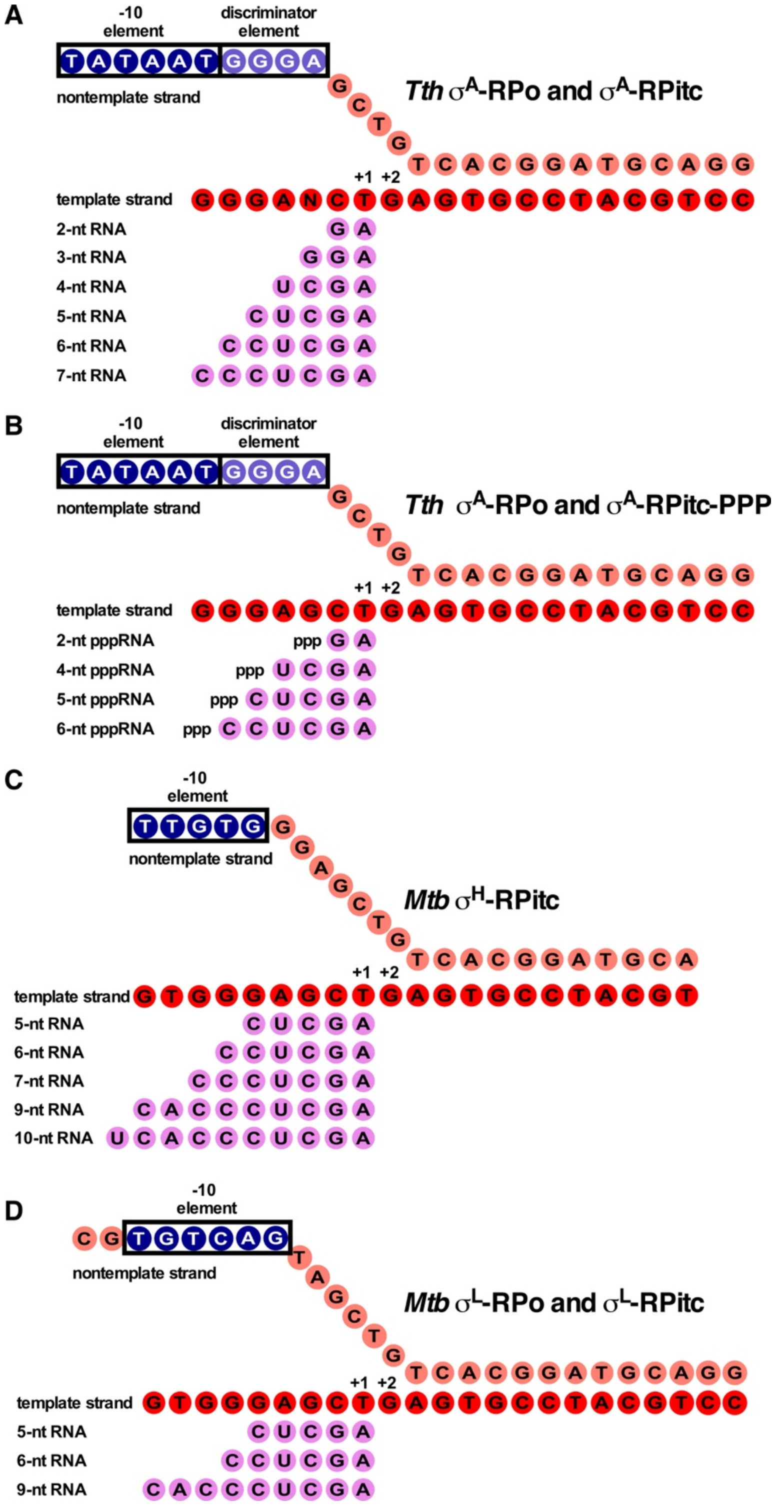


**Figure S1. Nucleic-acid scaffolds used for structural determination in this study.**

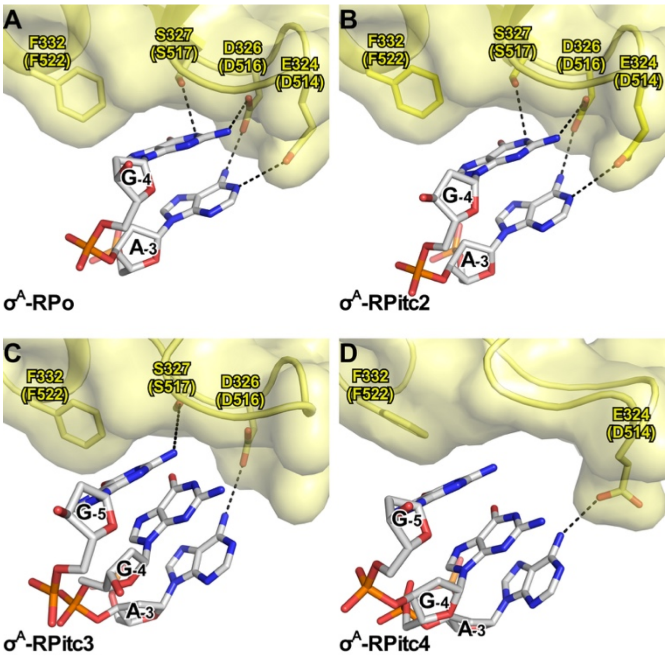


**Figure S2. Pre-organization by σ finger of the template-strand ssDNA.**
Interactions of σ finger with template-strand ssDNA in crystal structures of **(A)** *Tth* $\sigma^A$-RPo (PDB 4G7H) (ref. 1), **(B)** *Tth* $\sigma^A$-RPitc2 (PDB 4G7O) (ref. 1), **(C)** *Tth* $\sigma^A$-RPitc3 (this study), and **(D)** *Tth* $\sigma^A$-RPitc4 (this study). Yellow surfaces, solvent-accessible surfaces of σ; yellow ribbons, σ backbone; yellow and yellow-red stick representations, σ carbon and oxygen atoms; white, blue, red, and orange stick representations, DNA carbon, nitrogen, oxygen, and phosphorous atoms; black dashed lines, H-bonds.

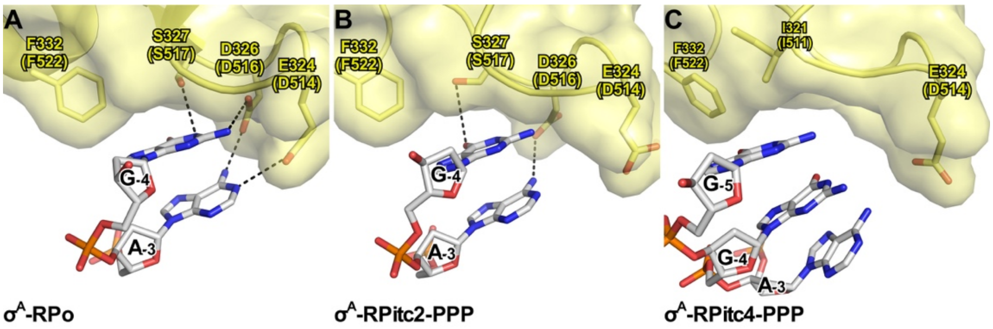


**Figure S3. Pre-organization by σ finger of the template-strand ssDNA.** The interaction of σ finger and template-strand ssDNA in crystal structure of **(A)** *Tth* $\sigma^A$-RPo (PDB 4G7H) (ref. 1), **(B)** *Tth* $\sigma^A$-RPitc2-PPP (this study), and **(C)** *Tth* $\sigma^A$-RPitc4-PPP (this study). Rendering and colors as in Fig. S2.

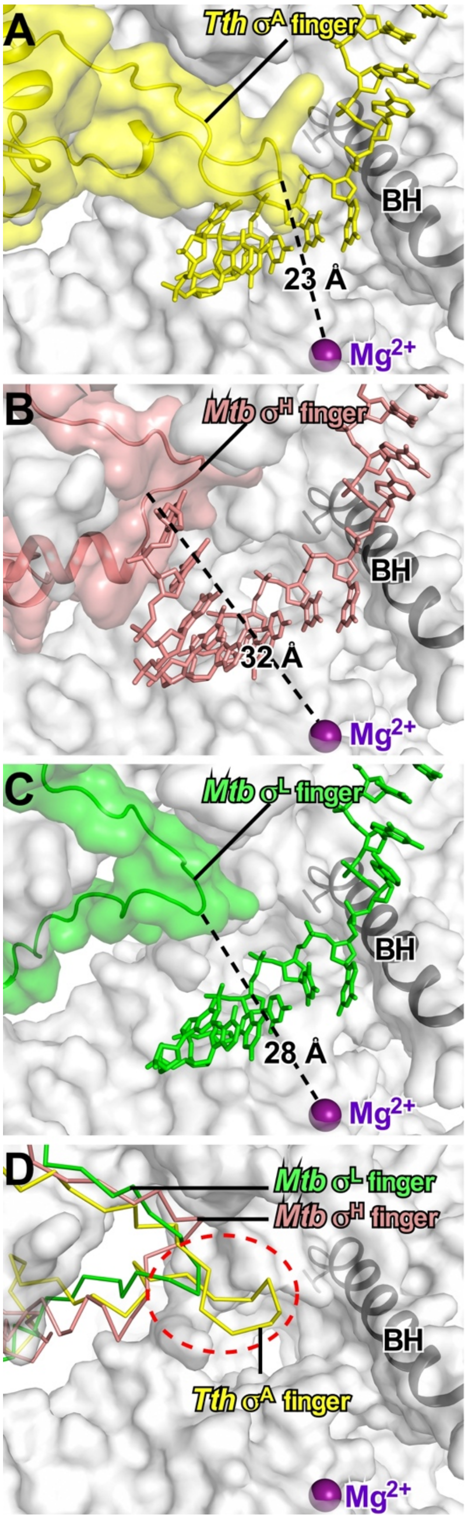


**Figure S4. RNAP active-center region and σ finger for primary σ factor, $\sigma^A$ (A), ECF σ factor $\sigma^H$ (B), ECF σ factor $\sigma^L$ (C), and superimposed $\sigma^A$, $\sigma^H$, and $\sigma^L$ (D).** RNAP active-center catalytic $Mg^{2+}$ ion, purple sphere; RNAP active-center bridge helix, gray ribbon; other RNAP residues, gray surface; σ finger, yellow, pink, and green ribbon plus surface for $\sigma^A$, $\sigma^H$, and $\sigma^L$, respectively (surfaces omitted in panel D); DNA template strand, yellow, pink, and green sticks for $\sigma^A$-, $\sigma^H$-, and $\sigma^L$-containing complexes, respectively (omitted in panel D). Distances between the tip of the σ finger and the RNAP active-center catalytic $Mg^{2+}$ ion, are indicated in panels A-C.

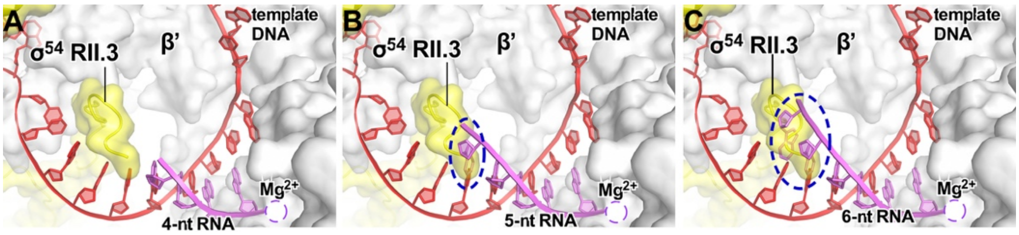


**Figure S5. Predicted stepwise displacement of transcription-initiation-factor module in bacterial $\sigma^{54}$-dependent transcription initiation; collision of RNA 5' end with $\sigma^{54}$ RII.3 upon extension of RNA from 4 nt to 5 nt.** Rendering and colors analogous to rendering and colors in Figs 1-4. Dashed ovals, predicted steric clashes.

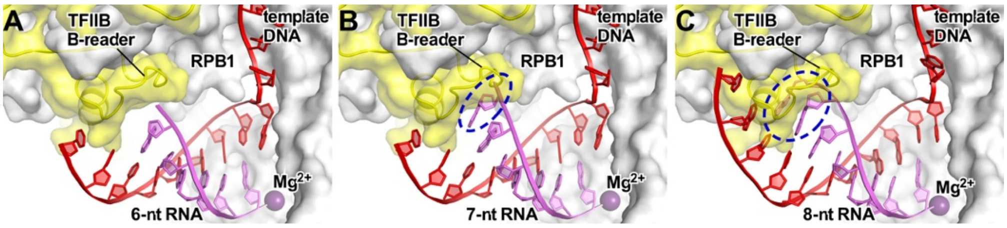


**Figure S6. Predicted displacement of transcription-factor module in eukaryotic RNAP-II-dependent transcription initiation: predicted collision of RNA 5' end with TFIIB B-reader upon extension of RNA from 6 nt to 7 nt.** Rendering and colors analogous to rendering and colors in Figs 1-4. Dashed ovals, predicted steric clashes.
